# Perfect representation of a convex 2D contour with only one Fourier component

**DOI:** 10.1101/2020.09.24.311316

**Authors:** Alf Harbitz

## Abstract

Shape analysis of a closed 2D contour is an important topic within biological shape analysis, where Fourier methods to reproduce the shape with a limited number of parameters have been and still are of vital importance. An example is within marine management research on fish, where shape analysis of otolith (earstone) contours is performed for species identification as well as for stock discrimination purposes. In both cases, it is expected that the fewer parameters that are needed in a method to reproduce the contour sufficiently good, the better. This contribution outlines how a convex contour of any shape can be represented to any wanted accuracy by only one Fourier component. The key idea is to allow a flexible choice of a predetermined number of *x*-values along an *x*-axis that goes through the two most distant points of the contour. The *y*-variable along the perpendicular *y*-axis is then monotonically transformed to a *z*-variable so that the minium and maximum *z*-values on the contour have the same distance from the *x*-axis. The *x*-values of the contour points are now chosen so that the corresponding *z*-values on the contour follows a perfect sinusoid if the *x*-values were equidistant. The method is illustrated by application to lasso contours of Norwegian Coastal Cod (NCC) and North East Arctic Cod (NEAC) otolith images, where the average new *x*-positions for the individual otolith contours were applied to all otoliths. The results show that a considerably better fit to the original individual otolith contours were obtained by applying the invers FFT to the new *y*-values than by the frequently applied 2D EFDs (Elliptical Fourier Descriptors) approach, for the same number, *m* < 11, of frequency components. A promising classification result was also obtained by the linear Fisher discrimination method and cross validation applied to the individual *x*-values for the NCC and NEAC otoliths, with 82% score for NCC and 80% score for NEAC with sample sizes 367 and 240, respectively.

## INTRODUCTION

Shape analysis of fish otolith contours for species identification, fish stock discrimination and other fishery management issues has been one of many fields with a rapidly increasing activity within biological shape analysis. Though there exist several methods for such analyses, Fourier methods [1] have been and still are the major approach in terms of scientific work reported in peer-reviewed journals. Since Fourier methods handle periodic phenomena, this is a natural approach to the analysis of a closed, 2-dimensional (2D) contour as we can think of a cursor tracing the contour in e.g. an anticlockwise direction several times providing a cyclic record with a smooth joint between each cycle. This is contrary to 1D time series records that normally will have an abrupt transition if repeated.

Among Fourier methods applied to shape analysis of a closed contour, Elliptical Fourier Descriptors (EFDs) [2], has been and is the dominant approach with an exponentially increasing number of scientific papers over the last decades [3]. The method is described in more detail in the method section, but briefly speaking it consists of approximating the contour with a superposition of ellipses with different tracing frequencies. Normally 10 frequencies will be sufficient to give a very good approximation to the large-scale structure of a 2D contour, even for rather complex shapes. Strangely enough the classical EFD needs more than one frequency component to represent a pure ellipse properly. In a recent paper [4] a modified EFD method was introduced that describes a pure ellipse perfectly with only one frequency component, and in addition appears to give better large scale approximations to general contours for a given number of frequency components than do the classical EFD.

A recipe for transforming a 2D contour to 1D is given in [5], which we denote MIRR because it mirrors the lower part of the contour around a vertical axis at the maximum *x*-value of the contour. The method is described in more detail in the method section. This approach applies FFT to the transformed *y*-coordinate values and needs half as many parameters per frequency component as EFD, but ambiguities easily occur for non-convex shapes. This method also reproduces badly a pure ellipse with only one frequency, but a modified method [4] based on a particular choice of non-equidistant *x*-values reproduces a pure ellipse virtually perfect with only one frequency component. In addition, the modified method appears to give better fit to the large-scale features of the original contour than the original method with the same number of frequency components.

Another popular Fourier method [6] uses the tangent angle to the contour as a basic variable instead of Cartesian coordinates as used by EFDs and MIRR. The continuous contour is created by a smooth interpolation between succeeding sampled contour points and a set of contour points is created with equidistant spacing along the contour. As for MIRR only two Fourier coefficients are needed for each frequency component and these are independent of each other. This approach also suffers from a good reproduction of a pure ellipse based on one frequency component, though it performs much better than the original EFD and MIRR in this case.

All the mentioned Fourier methods are aimed at describing periodic phenomena, and thus are not particularly appropriate to describe enhanced features of a contour on a very local scale. In the latter case, Wavelet transform has proven to be a useful alternative to Fourier methods [3],[7].

The main objective of this paper is to introduce a Fourier method, which we denote 1FC, which can reproduce any convex shape of a contour to any degree of accuracy with only one frequency component, by allowing a flexible choice of *x*-values adapted to the single contour under consideration. For non-convex shapes, the easily provided lasso contour coordinates [8] represent a guaranteed representation of the contour in terms of a convex polygon of lasso contour coordinates. The method is applied to samples of 2D contours of whole otoliths (earstones) extracted from images of Norwegian Coastal Cod (NCC) and North East Arctic Cod (NEAC) otoliths, where EFDs have proven to give good classification [9]. An example of a lasso contour of a whole NCC otolith is shown in Fig. 1. A direct discrimination approach with 1FC is applied by choosing a specific set of *x*-percentiles for each otolith as descriptors and by use of the linear Fisher discrimination method [10] combined with cross validation (leave one contour out at a time) to assess the method’s discrimination power. How well the 1FC method can reproduce an approximation to a contour from a known stock (e.g. NEAC) based on the average of the individual optimal *x*-values is examined by use of the FFT of the corresponding *y*-values for each contour. The goodness of fit is calculated based on the invers FFT for different numbers of frequency components and the results are compared with the classical EFD approach. It is also outlined how discrimination analysis can be performed based on the FFT of the individual *y*-values corresponding to the average of the characteristic average new *x*-values for each stock.

**Fig. 1.**
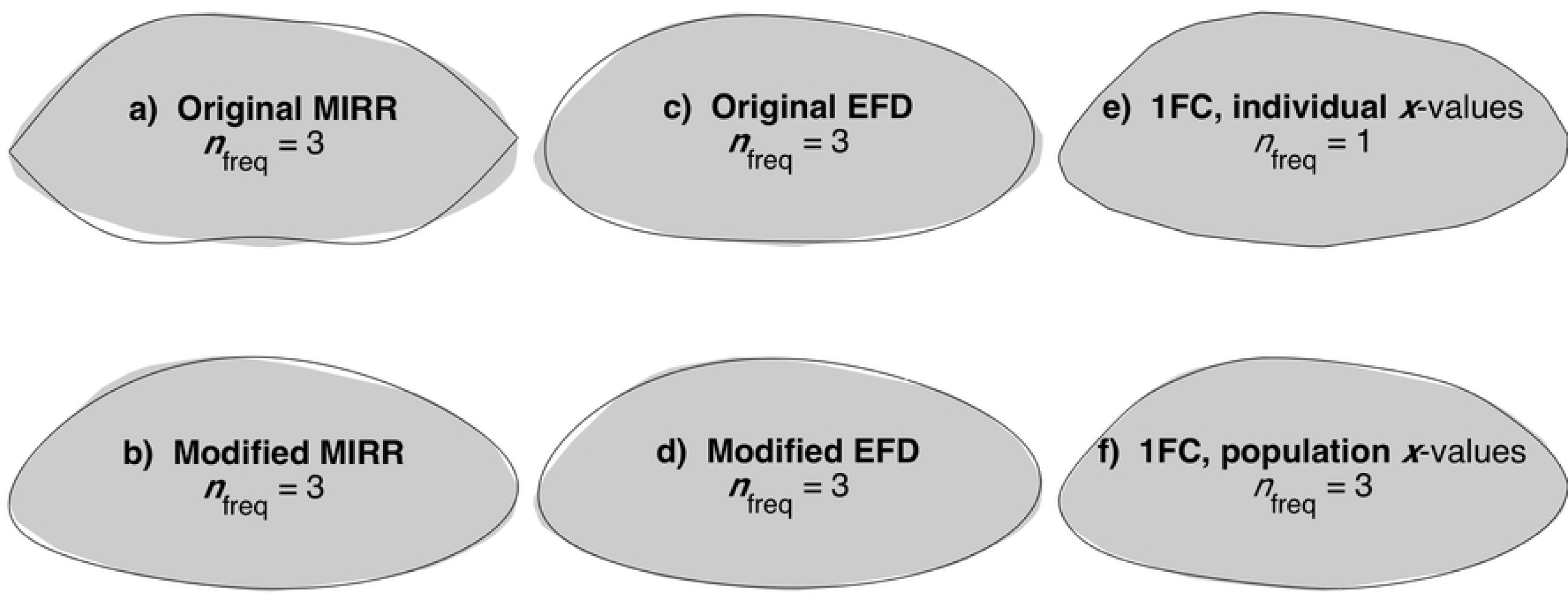
A typical NCC lasso contour with approximations by different Fourier methods with 3 frequency components except in e): a) Use of the original MIRR method, clearly showing false ringing features that disappear with the modified MIRR in b). In d) we also see a clearly better fit by the modified EFD compared to the result with the original EFD in c). In e) we see that by 1FC one frequency component is sufficient to give a virtually perfect reproduction of the original contour, and also a very good approximation by using the population x-values as seen in f).

## MATERIAL AND METHODS

### MATERIAL

The method outlined, along with other Fourier methods to be compared with, were applied to samples of NCC (367 otolith contours from the northern part of the Norwegian coast) and NEAC (240 otolith contours from the south-western part of the Barents Sea area). Stock discrimination results based on classical EFD methods are reported for the comparisons of NCC and NEAC in [9], which also reports discrimination results based on genetics for the same stocks, giving clear indication of differences between the stocks. In order to avoid false discrimination because of differences in fish length, the fish samples were restricted to fish lengths between 30 and 70 cm.

### FOURIER METHODS

#### Elliptical Fourier Descriptors (EFDs)

The EFD approach consists of approximating the contour with a superposition of ellipse trace vectors that trace each ellipse in a (normally) counterclockwise direction with different frequencies. The original EFD method is outlined in [2]. A nice feature of EFD is that there is no restriction on how the contour points are chosen. Standardization of the shape is found based on the first EFD ellipse. The original contour is rotated so that the major axis of the ellipse is horizontal, and each EFD is divided by the length of the major ellipse axis. Further, the order of the contour points are shifted so that the first point is the one closest to the rightmost point on the (rotated) first EFD ellipse. Each frequency component consists of 4 parameters, two characterizing the *x*-values and two characterizing the *y*-values. For the standardized EFDs, 3 of the 4 EFDs for the first ellipse are identical for all contours, while the fourth is the ratio between the minor and major axis of the first EFD ellipse.

A disadvantage of EFD is that the four descriptors for each ellipse are dependent, contrary to the two parameters describing a pure sinusoid in one dimension. Another property to be aware of is that the EFD approach is very sensitive to smoothing. In fact, small scale variations that normally are not visible without zooming may have a dramatic effect on the low frequency (large scale) EFD components. This sensitivity may easily lead to pitfalls in discrimination analysis providing false discrimination results. A method to examine if such pitfalls are present is outlined in [8]. In general smoothing of the contour is strongly recommended [6].

The reason why EFD performs badly on a pure ellipse is that the outline of the descriptors is based on the concept of a moving cursor with constant speed. This is in conflict with the classical mathematical description of an ellipse with *x* = *a* · cos*t* and *y* = *b* · sin *t* with time *t* as parameter, with speed *v* = *ds*/*dt* that is not constant unless for a circle where *a* = *b*. A modified EFD method using the appropriate tracing speed is given in [4] and demonstrates that a pure ellipse is now reproduced virtually perfectly by only one frequency component. In addition, the modified method appears to give a considerably better approximation to the large-scale feature of a fish otolith contour than the original EFD with the same number of frequency components up to 10 components, say.

#### The MIRR (partial reflection) Fourier method

We use the term MIRR (abbreviation for mirror) to denote the ‘partial reflection’ concept introduced in [5]. This is a method where the 2D contour is transformed to a 1D curve. Let *x*_min_ and *x*_max_ denote the minimum and maximum *x*-value of the horizontal coordinates of the contour points. The lower part of the contour between *x*_min_ and *x*_max_ is mirrored around a vertical axis through *x*_max_, and a set of new contour points (*x,y*) are calculated with equidistant *x*’s between *x*_min_ and *x*_max_. Then the FFT of the y-values are calculated, and a ‘large scale’ approximate reproduction of the 1D curve is found based on the invers FFT of the first frequency components. An advantage with this approach is that the FFT components are independent of each other with only two parameters (coefficients) per frequency component. To avoid ambiguities, however, the upper and lower parts of the contour need to be pure functions of *x*. This is the case for convex contours, which is the major focus of this paper.

For a pure ellipse with the *x*-axis along the major axis, the MIRR technique with equidistant *x*’s needs more than one frequency component to provide a good approximation, because the ellipse is vertical at *x*_min_ and *x*_max_. By replacing the equidistant *x*’s with flexible ones, however, a set of *x*-values can be chosen such that a pure ellipse will be virtually perfectly represented by only one frequency component [4]:

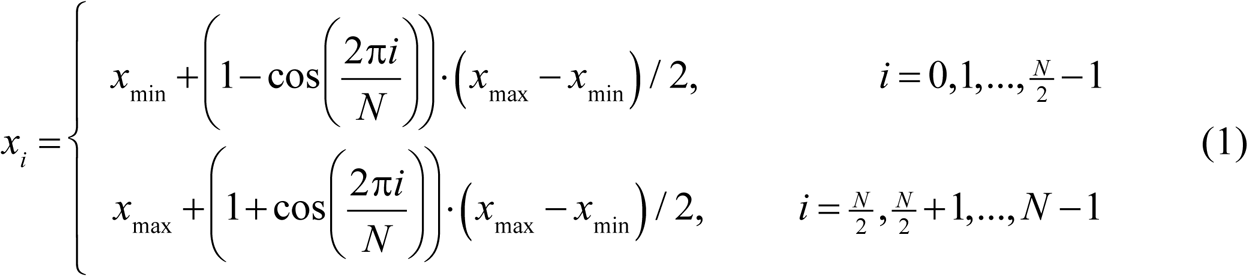

By the choice of *x*-values above it appears that MIRR applied to real fish otoliths gives better approximations to the large scale shape than does the MIRR with equidistant *x*’s, for the same number of frequency components (*ibid*).

#### The 1FC method

The 1FC method is a generalization of the modified MIRR method described above, and requires that the 2D contour can be transformed to a 1D curve described as a function of a horizontal *x*-variable without ambiguities. This requirement is satisfied for a convex contour, and in this case the transformed contour is found according to the following recipe (see Fig. 2):

**Fig. 2.**
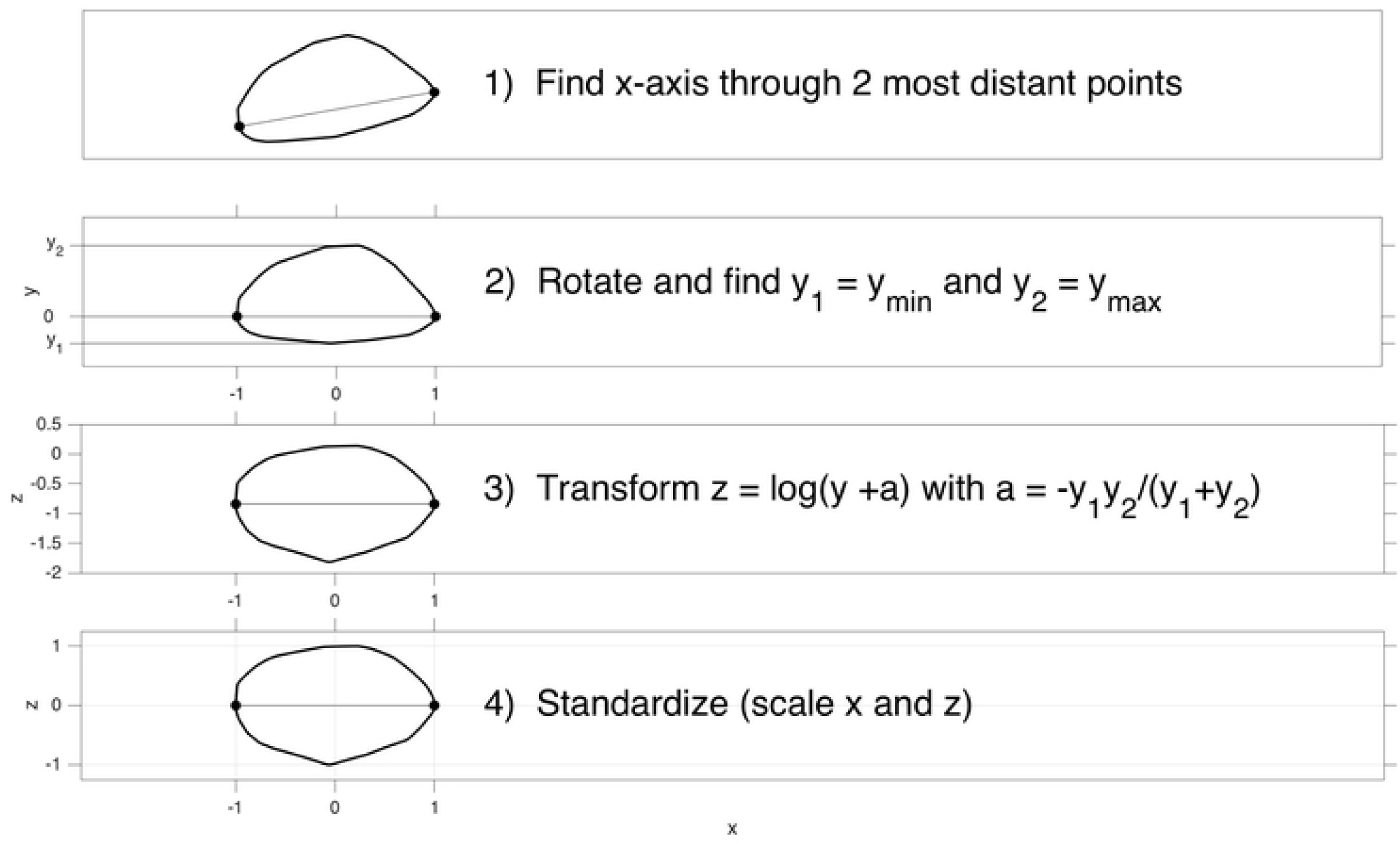
The 4-step procedure to produce the transformed contour coordinates for application of the 1FC method. Note that the transform in point 3) is one of a whole range of actual candidates, where a linear scaling of either the positive or the negative part in step 2) is the simplest approach.

1. Find the two most distant points on the contour and define the *x*-axis through these two points.
2. Rotate the contour so that the *x*-axis becomes horizontal and scale the *x*-variables so that *x*_min_ = −1 and *x*_max_ = 0. Define the *y*-axis perpendicular to the x-axis and let it be tangential to the transformed contour at *x*_max_.
3. Let *z* denote any monotonic transform of *y* so that *z*_max_ – *z*(*x*_max_) = z(*x*_max_) – *z*_min_, e.g. by the log-transform, *z* = log(*y*+*a*) with *a* = −*y*1*y*2/(*y*1+*y*2), as illustrated in Fig. 2, or by a linear scaling *z* = *y*/*y*max for *y* > 0 and *z* = *y*/*y*min for *y* < 0.

4.If *z*_max_ ≠ 1 or *z*_min_ ≠ –1 (as in the log-transform case), transform *z* linearly so that *z*_max_ = 1 and *z*_min_ = –1.

Note that the *a*-value in the log-transform in step 3 requires that *y*_max_ > –*y*_min_. If this is not the case, just set *y* = –*y* in step 3, and set *z* = –*z* after step 4.

When the original (*x,y*) are transformed to new (*x,z*) values as described above, the lower part of the contour is mirrored around the *y*-axis, so that *x*_max_ = 1. The concept for representation of the transformed contour to any wanted accuracy based on only one Fourier component is shown in Fig. 3. The black contour is the transformed original contour, and the mirrored lower part is shown by a dashed black curve. The red contour is a pure sinusoid where the red contour points (*x*_eq_,*y*_sin_) have equidistant *x*-values, *x*_eq_. We know from standard Fourier theory that for equidistant *x*-values a perfect representation of a pure sinusoid is straight forward as long as the period is an integer number of the distance *dx* between two succeeding *x*-values, and there are more than two points per period of the sinusoid (The Nyquist criterion, see [1]). The main idea is to find the same *y*_sin_-values on the transformed contour and the corresponding *x*_new_-values. Thus the FFT will only give contribution for the first Fourier component, and a perfect representation of the transformed contour is obtained by combining the inverse FFT with the new *x*_new_-values. Then the new points (*x*_new_,*y*_sin_) can be back transformed to (*x,y*) coordinates on the original contour by following the 4-step recipe for coordinate transformations backwards.

**Fig. 3.**
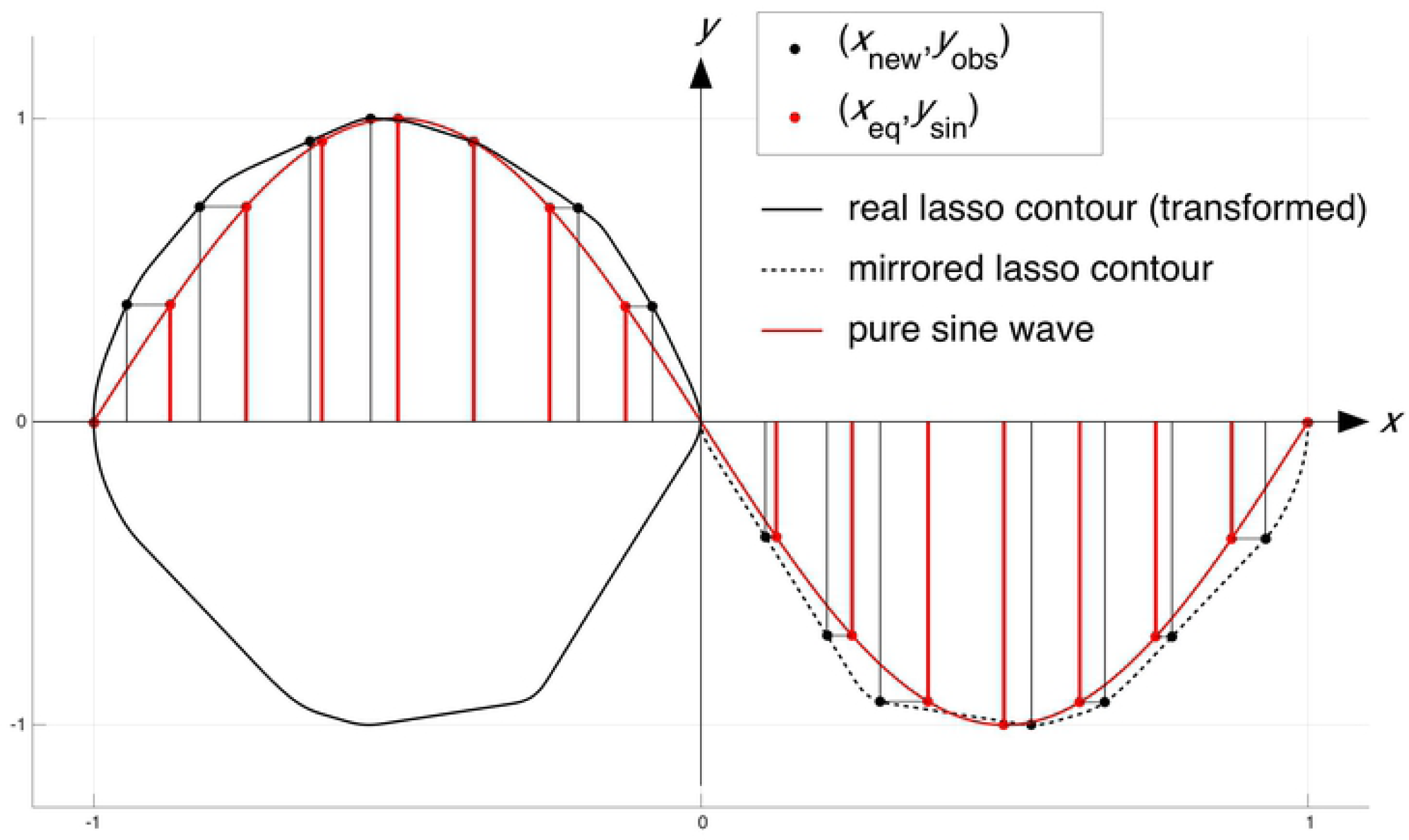
Illustration of how the new *x*-values by the 1FC method are found after the original convex contour is transformed as illustrated in Fig. 2. The red points that perform the pure sinusoid have equidistant *x*-values with a spacing *dx* so that the distance from *x*_min_ to *x*_max_ equals an integer multiplied by *dx*.

#### The 1FC is not restricted to convex contours

First of all, any non-convex closed contour can be represented by its convex envelope, or lasso contour, as illustrated in [8]. In addition, the recipe for the construction of 1FC only requires that the *y*-values after rotation of the original contour (Fig. 2) has only one maximum value and one minimum value. A biological example of a non-convex shape that satisfies this criterion is the tree leaf shown in Fig. 4. The green contour is based on back transformation from the 1FC approach with the scaled *y*-transform and based on only *m* = 256 coordinate points in the reproduction of the original contour with 1345 points. Note that in this case we see many deviations from a convex shape.

**Fig. 4.**
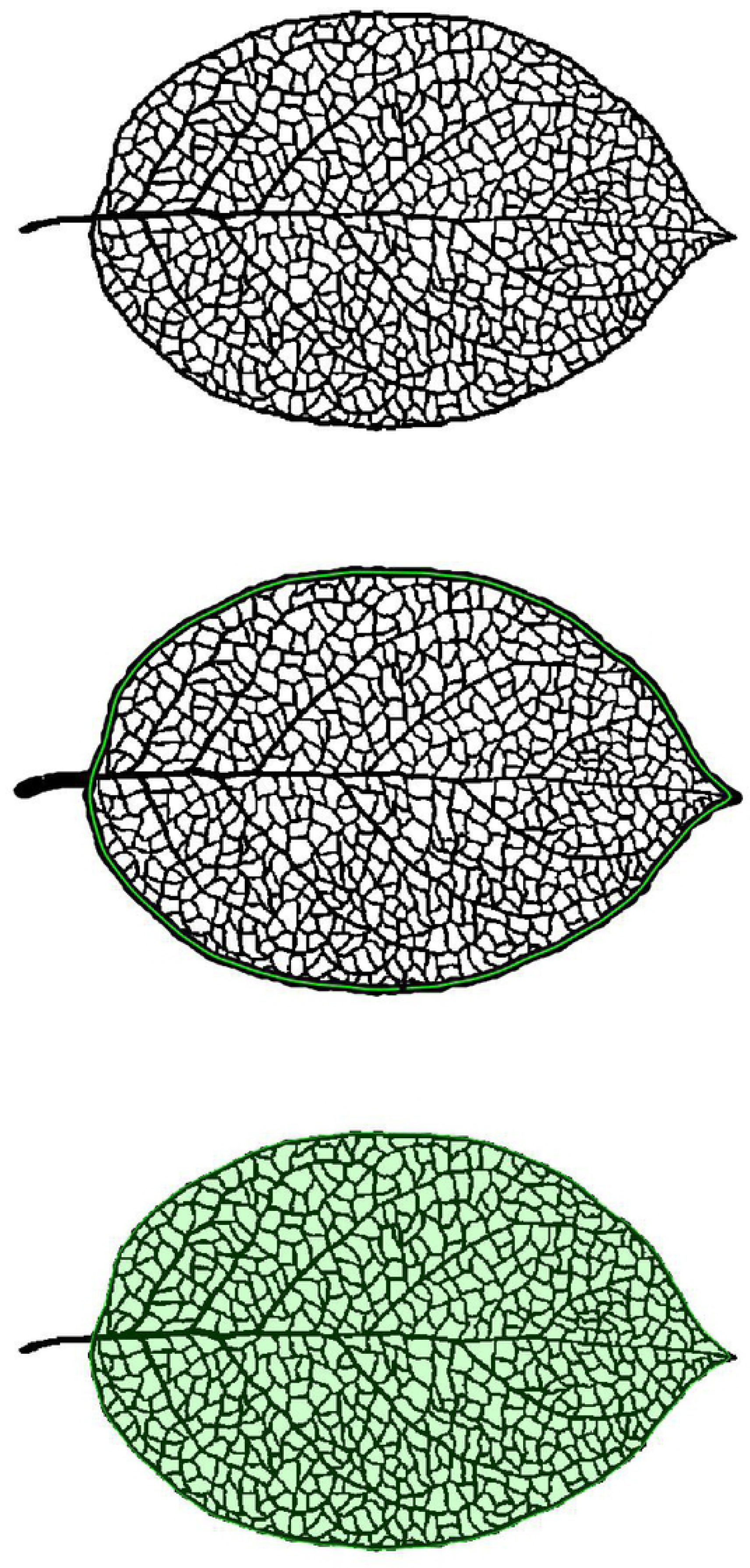
Illustration of a leaf with a non-convex contour, where the 1FC method can be applied. First the original contour is smoothed so that no local maxima or minima values of *y* are present, just one global maximum and minimum value. The upper panel shows the original contour. The mid panel shows the contour by a thick black line, with the 1FC reproduced contour in green with 256 x-values. The lower panel shows the original image with the green patched area defined by the 1FC coordinates.

#### The 1FC applied to a population of contours

A convex contour can be represented to any wanted accuracy with only one Fourier component by the 1FC with a sufficiently large number *n* of *x*_new_ values. How large *n* that is needed will depend on the shape of the contour as well as the location of the original (*x,y*) points on the contour. Some extra variables are also needed for the back transform process. So to assess if the 1FC method is useful as a data-reduction technique for individual contours as compared to other techniques like the use of a chain code [2], EFD or other, has to be done by experiments.

Say, however, that a perfect reproduction is not needed, but rather a decent reproduction that is sufficient to give the main characteristics for a population under study. A population in this context could for example be the sample of all otolith contours for a specific species. For the 1FC we imagine that a random sample of *n* individual contours are analyzed providing *n* distinct samples of standardized *x*_new_-values. Assume we have decided to construct *m x*_new_-values for each contour, *m* = 2048 being a typical example. We let the average of these *m* values, taken over all inidividual contours, be an estimated representation for the population under study.

Once a set of characteristic *x*_new_-values are created, we can examine how efficiently the individual contours can be represented based on this. First the *y*-values on the contour that correspond to the new *x*-values are calculated, then an FFT is applied to these new *y*-values. Thereafter, an inverse FFT is applied to the FFT with only a given number, *k*, of components, and the rest set to zero. Finally, the inverse FFT is back transformed to the original contour coordinates, and we measure how well the original contour is being represented as a function of the number of frequency components involved. Such results can be provided for different transforms within the 1FC method (e.g. the log- and linear transforms) as well as with other Fourier methods, to find the best method. Here we limit the attention to the comparison between the linear and log transform within 1FC, and comparison with EFD.

#### The application of 1FC in discriminant and classification analysis

Say that we want to apply the shape of the outer contour of a fish species, or a fish stock to find a measure to separate two populations, e.g. contours belonging to two different species of fish, or two different stocks of one species. We will consider two approaches, where the first and simplest one is to use a set of ranked x-values among the new ones provided by the 1FC method, as descriptors. As an example, let *x*_new(1)_ <*x*_new(2)_<…<*x*_new(n)_ denote the new *x*-values in increasing order. As a typical example let *n* = 2048, and apply each 20^th^ value: *x*_new(24)_, *x*_new(64)_, *x*_new(84)_,…, *x*_new(2024)_ as the descriptors for each otolith. If we now have *n*_1_ descriptors from one group with known identity, and *n*_2_ from another, we can for example apply Fisher’s linear discrimination method [10] directly and use cross validation (leave one out at a time) to examine how well these *x*-descriptors classify each otolith. We have no general recipe for how to choose the optimum number of *x*-predictors, and their corresponding order/percentiles, so this must be done by trial and failure. Some ideas might be created by comparing the average *x*-values for the two groups.

The other discrimination approach is to find the average of the two averaged *x*-values for each group, apply them to each otolith and find the corresponding *y*-values on the contour, and calculate the FFT of these new *y*-values. Fisher’s linear discrimination method can then be applied on a limited number of frequency components from the FFT and give an indication of discrimination success.

## THE QUANTIFICATION OF ACCURACY FOR A CONTOUR APPROXIMATION

The data to be used are the original contour coordinates (*x*_org_,*y*_org_) and the set of m points (*x*_fit_,*y*_fit_) on the approximate contour. As a measure of accuracy we calculate the mean distance between each (*x*_fit_,*y*_fit_) point and the closest point, (*x*_orgj_,*y*_orgj_), on the original contour, i.e. the set of continuous lines between all succeeding original coordinate points:

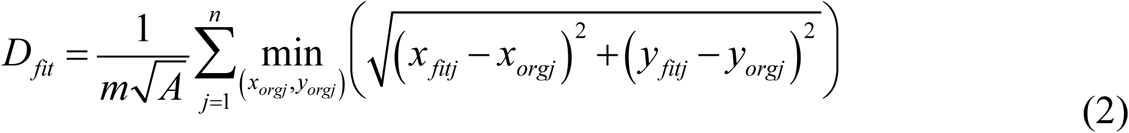

where *A* is the area enclosed by the closed contour and its role in the equation above is to make the fitness measure invariant to unit (or scale). The role of contour points and fitness points can easily be changed, and should give very similar results. If not, this indicates that the resolution is too coarse. As a measure of method goodness we apply the average *D*_fit_ value over all contours belonging to the group studied, e.g. the NCC sample.

## THE PROPORTION ESTIMATOR

Closely related to the classification of a group identity for an individual contour is the estimator for the proportion of the numbers belonging to the group relative to all other similar groups of interest. In the cod example studied it is particularly important to estimate the number of NCC, because this is considered to be a vulnerable stock. To do this one has to separate NCC from NEAC, which is done subjectively by visual otolith inspection. In principle this could be done automatically by shape analysis of otolith images. The direct estimate based on the detected fraction of NCC will be biased, however, unless the probability of correct classification is 100% for either stock. If we know the correct classification rates, a close to unbiased estimator for the unknown proportion *p* is as follows [11]:

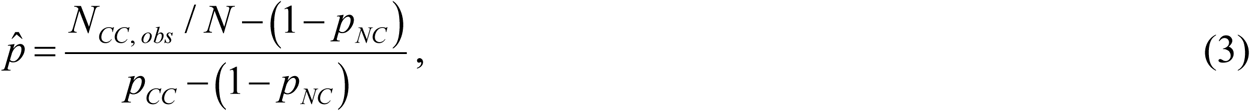

where *N*_*CC, obs*_ is the number of classified NCC, and N is the total number of NCC and NEAC. Further, *p*_*NC*_ is the probability of correct classification of a NEAC individual and *p*_*CC*_ is the probability of correct classification of a CC individual. This estimator is further discussed in the Conclusion and Discussion section.

## RESULTS

### GOODNESS OF FIT

The described 1FC method was applied to the sample of *n*_1_ = 367 NCC otolith lasso contours and *n*_2_ = 240 NEAC otolith lasso contours from images of the whole otolith, where *m* = 2048 standardized *x*-values were calculated for each otolith giving a virtually perfect reproduction of each otolith based on only one Fourier component. Then the average *x*-values for each of the two groups were applied and the corresponding *y*-values on each otolith contour in the group were calculated.

Fig. 5 shows the goodness of fit results by applying the averaged *x*-values and the approximate *y*-values based on the invers FFT of the new *y*-values on each otolith, as a function of the number of frequency components. Further, the results are calculated for both the log-tranform and the linear transform within the 1FC method, as well as by the classical EFD method.

**Fig. 5.**
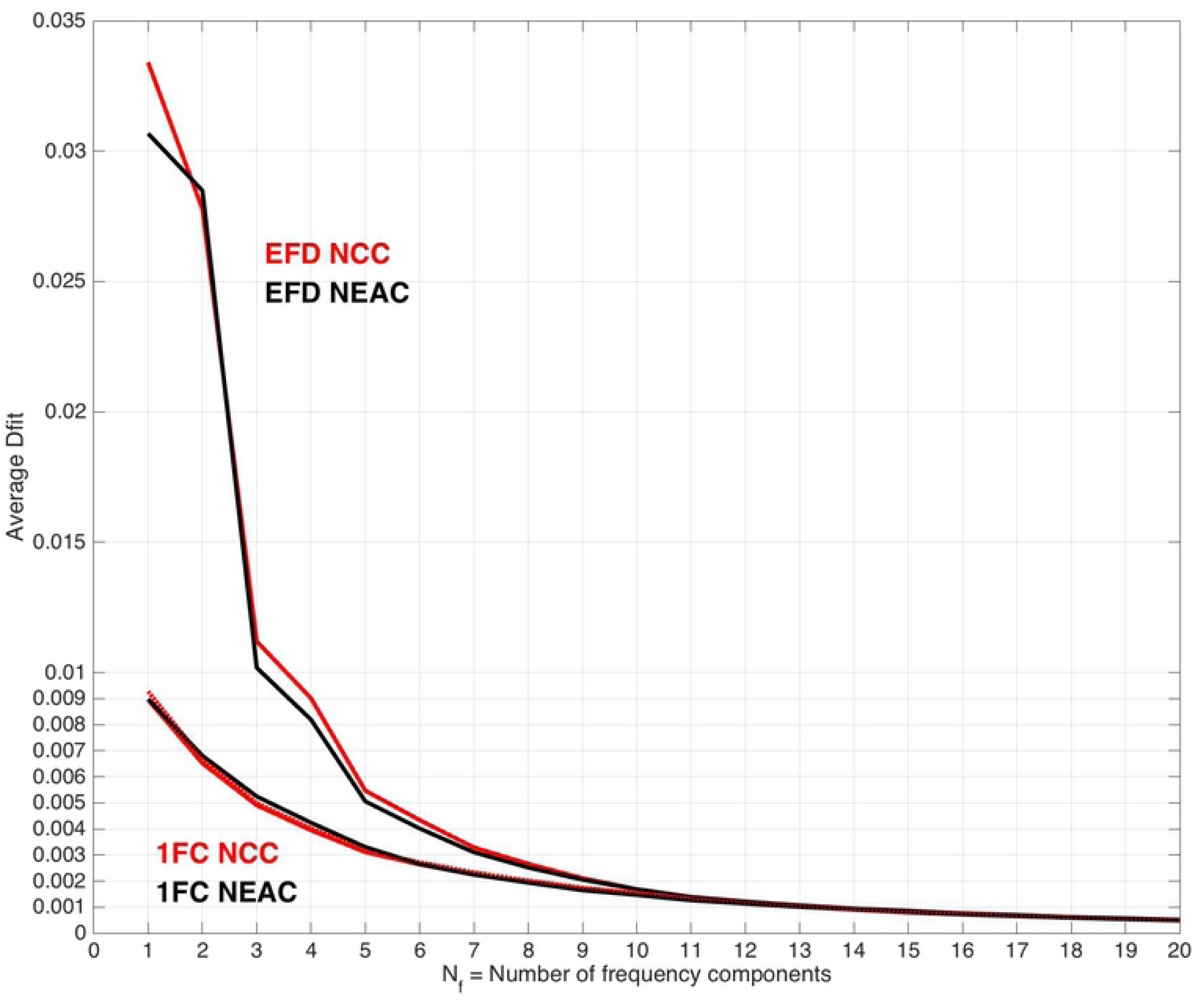
Results of 1FC reproduction of 367 NCC and 240 NEAC contours based on the average *x*_new_ values for each stock, compared to the EFD results (upper two curves). We see that the 1FC provides better results than EFD for the number of frequency components up to 10. The differences between the different 1FC results are hardly visible (log and linear *y*-transforms).

As we see from Fig. 5, the 1FC method performs much better than EFD with few parameters, and remain better until *n*_freq_ = 10. For larger *n*_freq_ values it is hard to see any difference between the methods. Though there are small differences between the 1FC approaches, the linear transformation of *y*-values give better results than the log-transformation, and the goodness of fit values are best for NCC for the *n*_freq_ values (2-5) where the difference between NCC and NEAC is largest.

### CLASSIFICATION OF NCC AND NEAC

By some trial and failure, the best classification of NCC and NEAC by Fisher’s linear discrimination and cross validation, based on the new *x*-values (m = 2048), was found for the 52 ordered values *x*_*new*(2:2:10)_, *x*_*new*(74:50:2024)_, *x*_*new*(2037:2:2047)_ with a probability of 82.83% correct score for NCC and 80.83% correct score for NEAC.

When the average of the average new *x*-values for NCC and NEAC were applied, the scores were 81.74% and 77.92% for NCC and NEAC, respectively, based on the 9 first frequency components. When adding the ratio between the max and min *y*-value with the x-axis through the most distant contour points, along with the ratio between the maximum extension in the *y*- and the x-direction, as descriptors, the scores increased to 85.56% and 82.50% for NCC and NEAC, respectively. These figures are not far from the scores 88.56% and 82.92% obtained by the classic EFDs based on 10 frequency components. The results based on 1000 bootstrap simulations are shown in Fig. 6. The average scores based on the data from which the bootstrap samples are drawn are shown as white vertical lines, and indicate some bias in some of the cases.

**Fig. 6.**
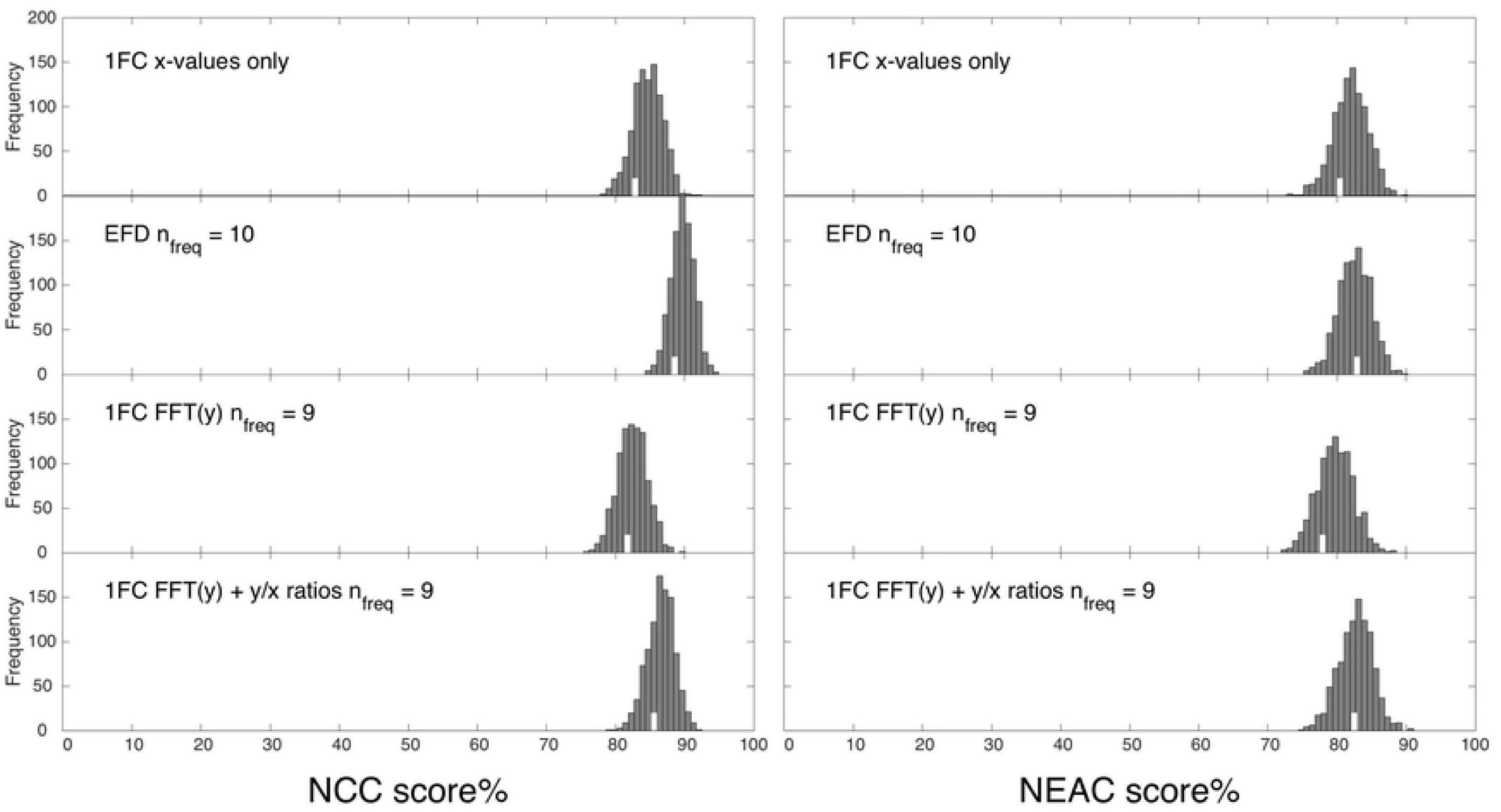
Discrimination results with 1FC and EFD applied to 367 NCC and 240 NEAC otolith contours based on 1000 bootstrap simulations. The *y*-values where FFT(*y*) is applied are the *y*-values on the transformed contour for the *x*-values being the average of the mean NCC and NEAC new *x*-values. The y/x ratios are the two ratio measures based on *y*_max_/*y*_min_ as well as the (*y*_max_-*y*_min_)/(*x*_max_-*x*_min_) values. The white vertical bars show the scores from data, which indicate biased values in some cases.

## CONCLUSION AND DISCUSSION

An easily applicable Fourier method, 1FC, is outlined that reproduces a contour described by a convex polygon of coordinate points to any wanted accuracy based on only one frequency component. The basic idea is to allow a flexible choice of coordinate x-values optimally adapted to the individual contour under consideration. To compare the performance of this method to reproduce the contour with other methods, like EFD, the average of the optimal *x*-values adapted to each contour in a particular group to be studied is applied to each contour in the group. Then an FFT is applied to the corresponding contour *y*-values, and an approximate reproduction of the contour is provided by back-transformation of the inverse FFT of the transformed *y*-values of the contour points. The results applied to the lasso contours of whole NCC and NEAC otoliths give far better approximations to the original lasso contours by 1FC than by EFD for the same number of frequency components until about 10, say, then the results merge.

Despite the much better fit to the large scale features of a contour that was obtained by 1FC, the discrimination results did not appear to be better than by EFD, though they were optimistically high. Because of the standardization of both the *x*- and *y*-coordinates by the 1FC method, possible shape differences characterized by different ratios between *y*- and *x*-values after rotation, but before scaling, are not taken appropriately into account. This was indicated by the substantial improvement based on the FFT of the transformed y-values when the two ratio-measures for the rotated (but not scaled) contour were added to the Fourier descriptors.

One would expect that a combination of methods with descriptors that detect different shape differences between two groups would have the potential to improve discrimination compared to the single method giving the best discrimination. Unfortunately such results were not obtained by combining the EFD descriptors and the 1FC descriptors applied to NCC and NEAC. An explanation to this can be that the two methods are not different enough with regard to the actual shape studied, and that the success of combining methods may vary greatly with shape.

It is a promising result that the set of *x*-values providing the “perfect” reproduction of a contour with only one frequency component could be used directly as a pool of descriptors that gave good discrimination results in our case. The results, however, were sensitive to the number of x-values that was chosen along with their order (percentile values). This might be an interesting field for more thorough studies.

Another field for further exploration is to study the estimator of the proportion of a group. Within fisheries management of for example Norwegian Coastal Cod, which is a vulnerable stock, it is important to detect the fraction of NCC in catches with cod. This is done by subjective visual inspection of the otoliths today, but can in principle be done by automatic image analysis of the whole otolith. If the correct classification rate is known for both NCC and NEAC, the ratio based on detected stocks will be biased and can be replaced by the close to unbiased estimator given by Eq. 3. If we have different methods, like EFD and 1FC, giving different, but close to unbiased results, these can be combined and hopefully reduce the variance of the estimator.

The 1FC method is based on an *x*-axis going through the most distant points of the original contour. In cases that this is not optimal one can play with a transformation of the coordinates so that the most distant points in the transformed space appear to be more appropriate. A simple example of this is a scaling of one of the two coordinates.

